# Census of exposed aggregation-prone regions in proteomes

**DOI:** 10.1101/2022.12.16.520802

**Authors:** Théo Falgarone, Etienne Villain, Francois Richard, Zarifa Osmanli, Andrey V. Kajava

**Affiliations:** Centre de Recherche en Biologie cellulaire de Montpellier, CNRS, Université Montpellier, Montpellier, 34293, France; Biophysics Institute, Ministry of Science and Education of Azerbaijan Republic, Az1141, Baku, Azerbaijan; Institut de Biologie Computationnelle, Université Montpellier, 34095 Montpellier, France

**Keywords:** protein aggregation, bioinformatics, large-scale analysis, evolution, kingdoms of life

## Abstract

Loss of solubility usually leads to the detrimental elimination of protein function. In some cases, the protein aggregation is also required for beneficial functions. Given the duality of this phenomenon, it remains a fundamental question how natural selection controls the aggregation. The exponential growth of genomic sequence data and recent progress with *in silico* predictors of the aggregation allows approaching this problem by a large-scale bioinformatics analysis. Most of the aggregation-prone regions are hidden within the 3D structures and, therefore, they cannot realize their potential to aggregate. Thus, the most realistic census of the aggregation prone regions requires crossing aggregation prediction with information about the location of the natively unfolded regions. This allows us to detect so-called “Exposed Aggregation-prone Regions” (EARs). Here, we analyzed the occurrence and distribution of the EARs in 76 full reference proteomes from the three kingdoms of life. For this purpose, we used a bioinformatics pipeline, which provides a consensual result based on several predictors of aggregation. Our analysis revealed a number of new statistically significant correlations about the presence of EARs in different organisms, their dependence on protein length, cellular localizations, co-occurrence with short linear motifs, and the level of protein expression. We also obtained a list of proteins with the conserved aggregation-prone sequences for further experimental tests. Insights gained from this work led to a deeper understanding of the functional and evolutionary relations of the protein aggregation.

## INTRODUCTION

Proteins are usually soluble molecules interacting transiently with each other or the other biomolecules. After performing their functions, they are degraded by proteases. Thanks to the dynamic balance between protein synthesis and degradation, living organisms can efficiently regulate many different processes. However, occasionally, some proteins, often for not entirely clear reasons, form aggregates. Most of the aggregates have a very characteristic structure of amyloid fibrils. These fibrils are typically straight, around 10 nm in diameter, thermostable, protease resistant, and rich in β-structure (1). They are completely or partially insoluble and frequently lead to a variety of age-related diseases including Alzheimer’s disease, Parkinson’s disease and others (2). In some cases, the amyloid fibrils (named prions) can be “infectious agents”. The prion fibrils, which are found themselves in another organism or a cell, can trigger the formation of similar fibrils and cause transmissible neurodegenerative diseases (3). The amyloid deposits can not only be composed of copies of the same protein, but also represent co-aggregates of two or more proteins and by doing so simultaneously impair several biological processes (4). At the same time, not all amyloid fibrils are linked to diseases. Increasing number of studies describe so called “functional” amyloids, which fulfill beneficial roles in the organism (5, 6). For example, curli proteins from some gram-negative bacteria form amyloid fibrils on the bacterial surface. They are involved in biofilm formations, which is a successful strategy allowing microorganisms to resist the threats of the environment (UV radiation, oxygen, desiccation etc) (7). Other examples from mammals are RIP1 and RIP3 proteins whose co-aggregation into amyloid fibrils mediates a key interaction of necroptosis signaling (8, 9).

Despite great interest in protein aggregation, especially regarding amyloids, scientists have focused on a few of the most devastating amyloidoses or known cases of functional amyloids. However, the overall prevalence of the protein aggregation in organisms is not yet well studied. This analysis requires computational methods for *in silico* prediction of the aggregation. The propensity to form aggregates is coded by the amino acid sequence, therefore, several computational programs have been developed (10–18) Availability of the computational tools for prediction of aggregation-prone regions made it possible to obtain a more general view of this phenomenon by using *in silico* analysis of the whole-proteome data. Previous *in silico* studies revealed a number of interesting observations (14, 19–28). For example, a study of six proteomes (*P. tetraurelia, S. cerevisiae, C. elegans, D. melanogaster, M. musculus* and *H. sapiens*) using a specially developed algorithm, demonstrated that the average aggregation propensity of a proteome correlates inversely with the complexity and longevity of the studied organisms (29). In another analysis of the proteomes of *D. melanogaster, S.cerevisiae* and *C. elegans* using TANGO predictor (13) it was shown that proteins that are essential to organism fitness (knockdown of these genes leads to lethality), have a lower aggregation score than nonessential proteins (23). Analysis of the human proteome by the Zyggregator method (30, 31) suggested that proteins involved in the secretion pathway are more prone to aggregation compared to non-membrane proteins in general (27). Application of the 3D profile method to *E. coli*, *S. cerevisiae*, and *H. sapiens* proteomes showed that the predicted high propensity for amyloid formation does not reflect well the limited number of proteins involved in disease-related or functional amyloid deposits (26). The same analysis of proteins from PDB suggested that most of the predicted aggregation prone regions are hidden within the 3D protein structure and, therefore, inaccessible for intermolecular interactions such as aggregation (32). The analysis of cytosolic bacterial (*E. coli*) and eukaryotic (*H. sapiens*) proteomes indicated that the aggregation propensity of proteins inversely-correlates with their abundance (19–21). Most of these data are in agreement with the conclusion that the evolutionary pressure acts on the proteins to minimize their aggregation propensity.

Several publications have reported that proteomes contain a very high percentage of proteins with amyloidogenic or aggregation-prone regions (AR), which is in obvious conflict with a small number of the known proteins involved in amyloidoses (28). It was explained by the fact that most of the predicted ARs are hidden within the 3D structure preventing aggregation (33). Conversely, in most known cases of amyloidosis, the native conformation of the polypeptide chains that form amyloid deposits *in vivo*, is unfolded (or intrinsically disordered). Thus, to get a more realistic census of the aggregation prone regions in proteomes, it is necessary to cross aggregation prediction with information about the location of the intrinsically disordered regions (IDRs). IDRs are always exposed for the intermolecular interactions critical for aggregation. We used this concept to develop a computational pipeline TAPASS (33), which can detect such “Exposed Amyloidogenic Regions” or, otherwise “Exposed Aggregation-prone Regions” (EARs) located within IDRs and carrying high potential to aggregate (see Figure 1). To obtain the most consensual results on the occurrence and distribution of the EARs in proteomes we selected three predictors of aggregation (TANGO, Pasta 2.0 and ArchCandy 2.0) (10, 13, 16). They were selected based on the diversity of their basic principles, their popularity, and ability to be downloaded for analysis of a large number of sequences. TAPASS also provides information about the cellular localization, post-translational modifications, and functions of aggregation-prone proteins.

**Figure 1.**
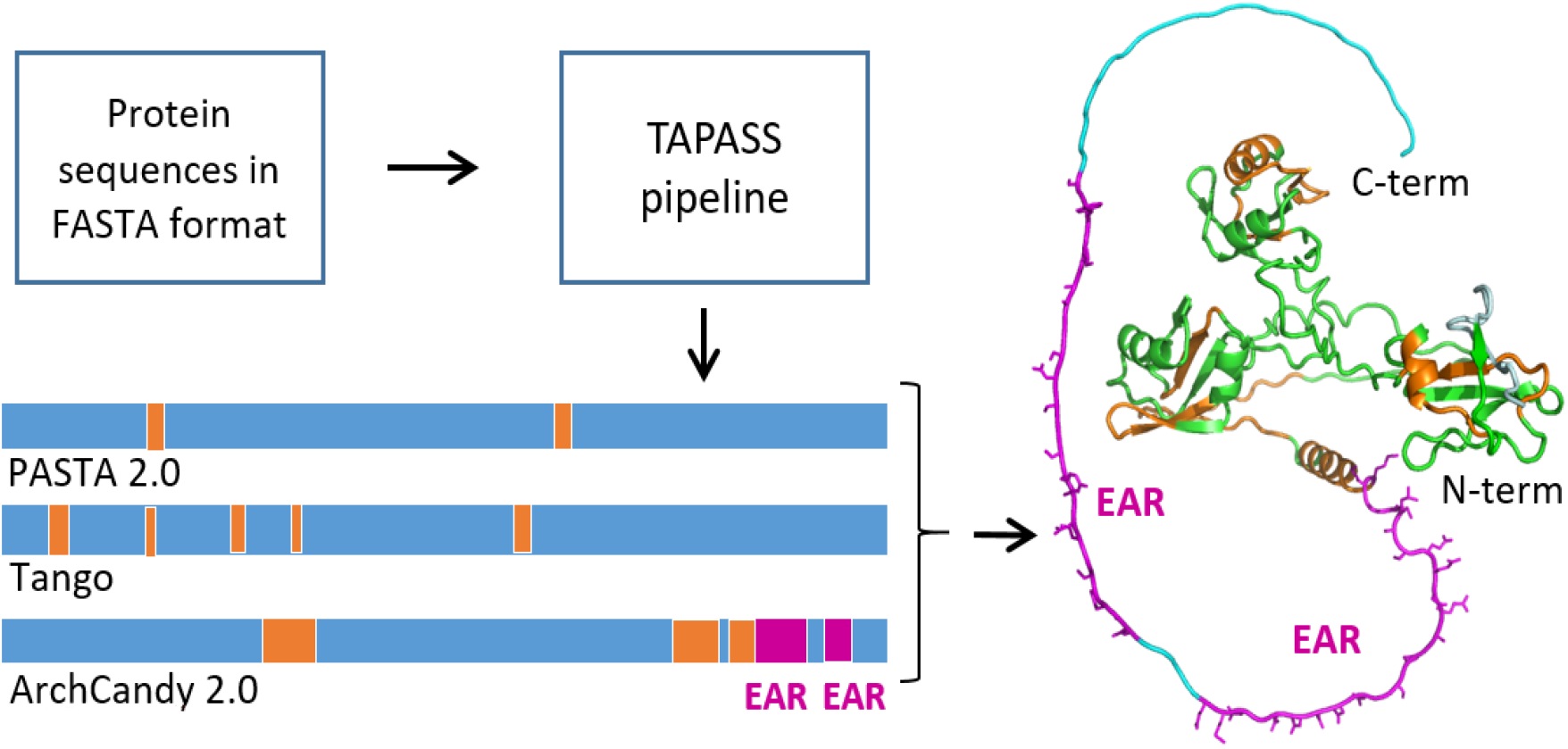
A general scheme showing mapping of ARs and EARs on a structural model of human TAR DNA-binding protein 43. This protein forms amyloid fibrils by the C-terminal Low Complexity Domain (LCD, 274-414) (34). TAPASS predicts several ARs, which are located within the 3D structures (orange) and two EARs (magenta) located at the C-terminal IDR.

In addition to the advances with the predictors of aggregation, the past few years were marked by a significant increase in the number and quality of proteome sequencing data, Thus, the advances with methods predicting aggregation potential, development of the TAPASS pipeline, as well as an increasing number of high quality whole-proteome sequencing data, made a new census of aggregation-prone regions in proteins timely. In this paper, we present the results of such a detailed analysis of 76 full reference proteomes from the UniProt databank.

## MATERIAL AND METHODS

### TAPASS pipeline

The input file of TAPASS requires protein sequences in Fasta format and can contain additional informations from UniProt (35) (gene id, GO term, version, modification date…). The pipeline uses: IUPred (36) and our in-house predictor (IDRs), CATH associated with HMMER 3.3 (structural domains)(37, 38), TMHMM (transmembrane regions) (39), SignalP (signal peptide) (40), SLiMs (short linear motifs) (41, 42), Pfam (structural and functional domains) (43), Pasta 2.0 (16), TANGO (13) and updated version of ArchCandy 2.0 (aggregation-prone regions) (10, 33).

The results of the three predictors of aggregation, ArchCandy 2.0, Pasta 2.0 and TANGO, were treated separately. Each predictor gives the start and end positions of ARs in protein sequences. An AR is considered as EAR if at least 80 % of an individual hit of AR predictor overlap with an IDR. Thus, our analysis led to three independent censuses of the aggregation-prone regions. If all three censuses yielded similar regularities, then these findings were considered as more reliable and treated with special attention.

### Selection of proteomes for large-scale analysis

76 reference proteomes with 1 123 749 proteins in total were selected from the UniProt databank (see Supplementary data) (35). The proteomes belong to the three kingdoms of life: eukaryote, bacteria and archaea. The selection of species was made to have well-annotated and complete reference proteomes covering the diversity of living organisms. Viral proteomes were not considered in this analysis due to small size of their proteomes yielding very different results depending on the strains. Their analysis will be a subject of our future study.

## RESULTS

### Occurrence of ARs and EARs in the proteomes

Previous studies detected a very high percentage of AR-containing proteins in proteomes with almost each protein having at least one predicted AR (22, 26, 27). The results of our analysis of 76 reference proteomes support this conclusion predicting 68.6 %, 79.3 % and 90.0 % of AR-containing proteins by ArchCandy 2.0, Pasta 2.0 and TANGO, respectively. The coverage of ARs, obtained by dividing the number of amino acid residues involved in ARs by the number of all residues in proteins, is equal to 12.6 %, 6.2 % and 11.3 % for ArchCandy 2.0, Pasta 2.0 and TANGO respectively. A very high percentage of AR-containing proteins is in contradiction with a small number of proteins known to be involved in different amyloidoses or functional amyloids. However, if we consider EARs, the number of potential aggregation-prone proteins is drastically reduced. EAR-containing proteins represent 9.0 %, 6.8 % and 19.5 % of all proteins with coverage of 0.8 %, 0.2 % and 0.4 % of residues according to ArchCandy 2.0, Pasta 2.0 and TANGO respectively. The low percentage of proteins with EARs, in contrast to a very high percentage of ARs, agrees better with the small number of the known proteins involved in aggregation *in vivo*.

### Aggregation-prone regions in prokaryotic and eukaryotic organisms

Analyzing the 76 selected proteomes we observed a relatively uniform distribution of AR-containing proteins among the organisms (Figure 2). Curiously, *Homo sapiens* has the least number of AR-containing proteins. At the same time, we saw a large variation in the proportion of EAR-containing proteins. Among the organisms with the least number of EAR-containing proteins are thermophilic prokaryotes (6 archaea and 5 bacteria: *Chloroflexus aurantiacus, Thermodesulfovibrio yellowstonii, Dictyoglomus turgidum, Nanoarchaeum equitans, Sulfolobus solfataricus, Thermotoga maritima, Archaeoglobus fulgidus, Thermococcus kodakaraensis, Methanocaldococcus jannaschii, Candidatus korarchaeum, Aquifex aeolicus*).

**Figure 2.**
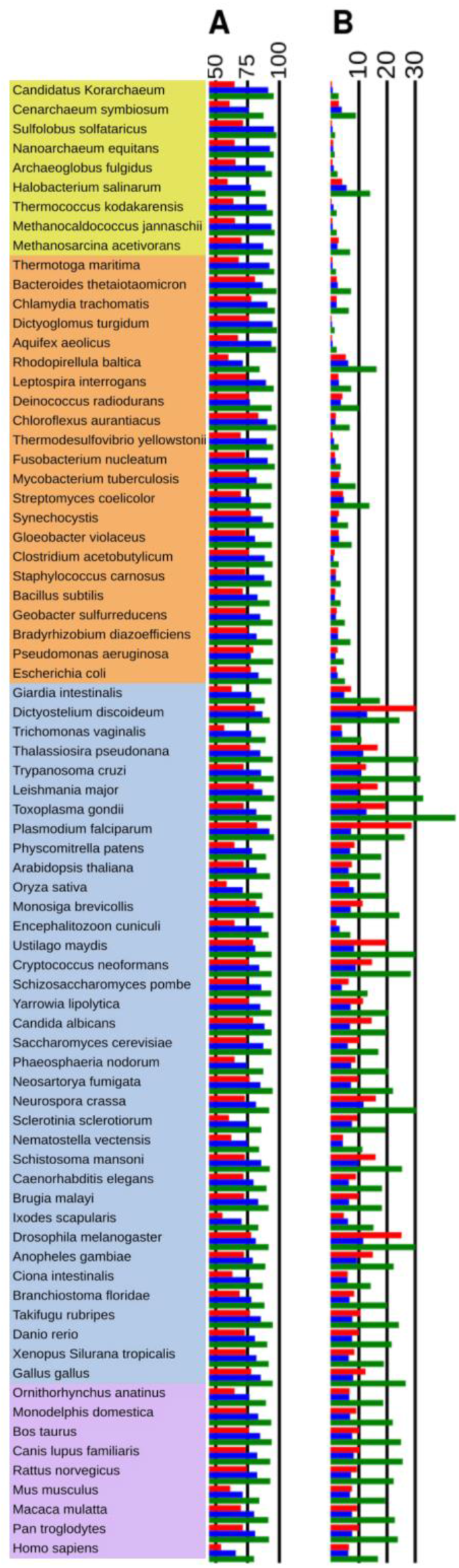
Proportion of (A) AR- and (B) EAR-containing proteins per organism predicted by using three predictors of aggregation, ArchCandy 2.0 (red), Pasta 2.0 (blue) and TANGO (green). Archaea, bacteria, eukaryotes and mammalian eukaryotes are outlined by yellow, orange, blue and violet respectively (made by using free options in iTOL, https://itol.embl.de/ (45)).

The eukaryotes with the simplest level of organization, mostly unicellular (or partially unicellular) protists such as *Plasmodium falciparum, Leishmania major, Thalassiosira pseudonana, Trypanosoma cruzi, Toxoplasma gondii and Dictyostelium discoideum* have the greatest numbers of EAR-containing proteins (Figure 2). High levels of EAR-containing proteins are also found in two fungi (*Ustilago maydis, Neurospora crassa*), fruit flies (*Drosophila melanogaster*), mosquitoes (*Anopheles gambiae*) and chickens (*Gallus gallus*). Most of them are known to have the greatest number of low-complexity repetitive sequences (44). This is particularly the case of *Trypanosoma cruzi* and *Dictyostelium discoideum*, which have an abnormal high level of Asn/Gln rich regions, two types of amino acids frequently found in amyloids. Among analyzed mammalians, *Homo sapiens* has the least number of EAR-containing proteins (Figure 2).

Having a global view of the dispersion of aggregation potential of the proteomes, it was interesting to analyze the tendencies associated with groups of the organisms. First, we compared prokaryotes and eukaryotes. All three predictors detect more AR-containing proteins and higher AR-coverage in prokaryotes in comparison to eukaryotes (Figure 3A and 3B). The tendency is reversed when we compare the occurrence of EARs (Figure 3C). The percentage of EAR-containing proteins and coverage of EARs are noticeably higher in eukaryotic than in prokaryotic organisms (Figure 3C and 3D). This can be explained by a higher number of IDRs in eukaryotes, which require the IDRs to mediate a more complex network of protein-protein interactions in comparison to prokaryotes (46, 47). At the same time, the coverage of EARs in IDRs is lower in eukaryotes compared to prokaryotes (Figure 3E). Thus, the eukaryotic IDRs are less aggregation-prone on average than the prokaryotic ones, suggesting a higher selective pressure on their IDRs to avoid aggregation.

**Figure 3.**
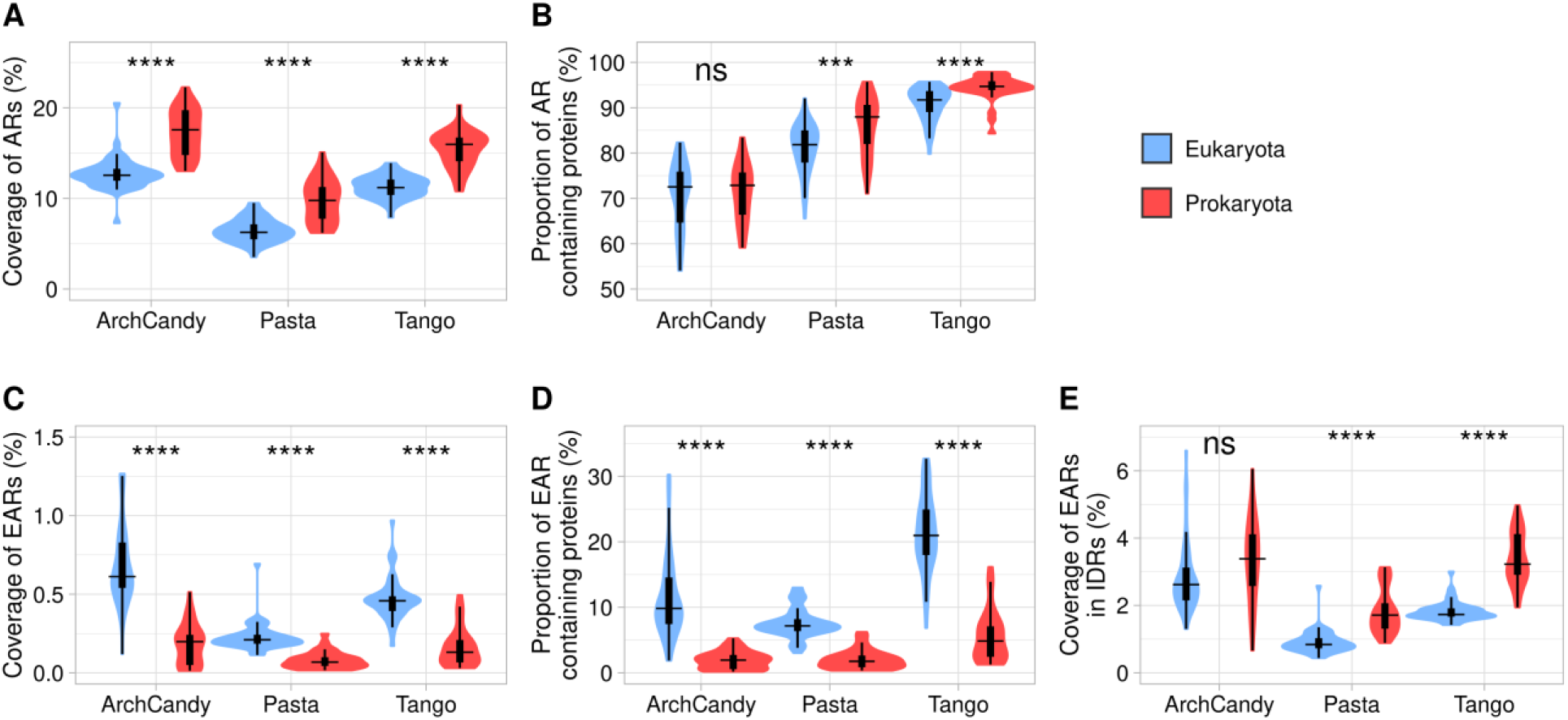
Level of aggregation potential according to three amyloid predictors in prokaryotes and eukaryotes. Coverage of ARs (A), proportion of AR-containing proteins (B), coverage of EARs (C), proportion of EAR-containing proteins (D) and coverage of EARs in IDRs (E). For statistical analysis between eukaryotic and prokaryotic organisms we performed a t-test for the predictors individually (ns: non-significant; *: p-value < 0.05; **: p-value < 0.01; ***: p-value < 0.001; ****: p-value < 0.0001).

### The more thermophilic the less aggregation-prone

A unique feature of prokaryotes is the wide range of their optimal growth temperatures (OGTs), some of them reaching temperatures above 105°C (48). We estimated the aggregation potential of the prokaryotic proteomes depending on the OGTs. For this purpose, we subdivided the selected reference proteomes into two groups: 20 mesophilic organisms, those with an OGT below 41°C and 11 thermophilic organisms with an OGT above 41°C. The comparison of proportion and coverage of ARs from these groups do not reach the same conclusion as ArchCandy 2.0 predicts a decrease in ARs in the thermophilic organisms, while PASTA 2.0 and TANGO show the opposite tendency (see Supplementary Figure 1). However, evaluation of EARs by all the predictors clearly demonstrated that they decrease with the increase of OGT (Figure 4). It has been also shown that the frequency of glutamine residue, which has a high amyloidogenic potential, decreases, while the total frequency of charged residues, which can block amyloid-formation, increases in thermophilic proteins(49). At the same time, the temperature increase may favor aggregation. For example, it has been shown that the amyloidogenesis rate constant of Aβ-peptide increases and the lag time decreases with increasing temperature (50). Considering all this, we can conclude that the decrease in the EARs with OGT can be a result of an evolutionary pressure on the thermophilic proteins to avoid the aggregation.

**Figure 4.**
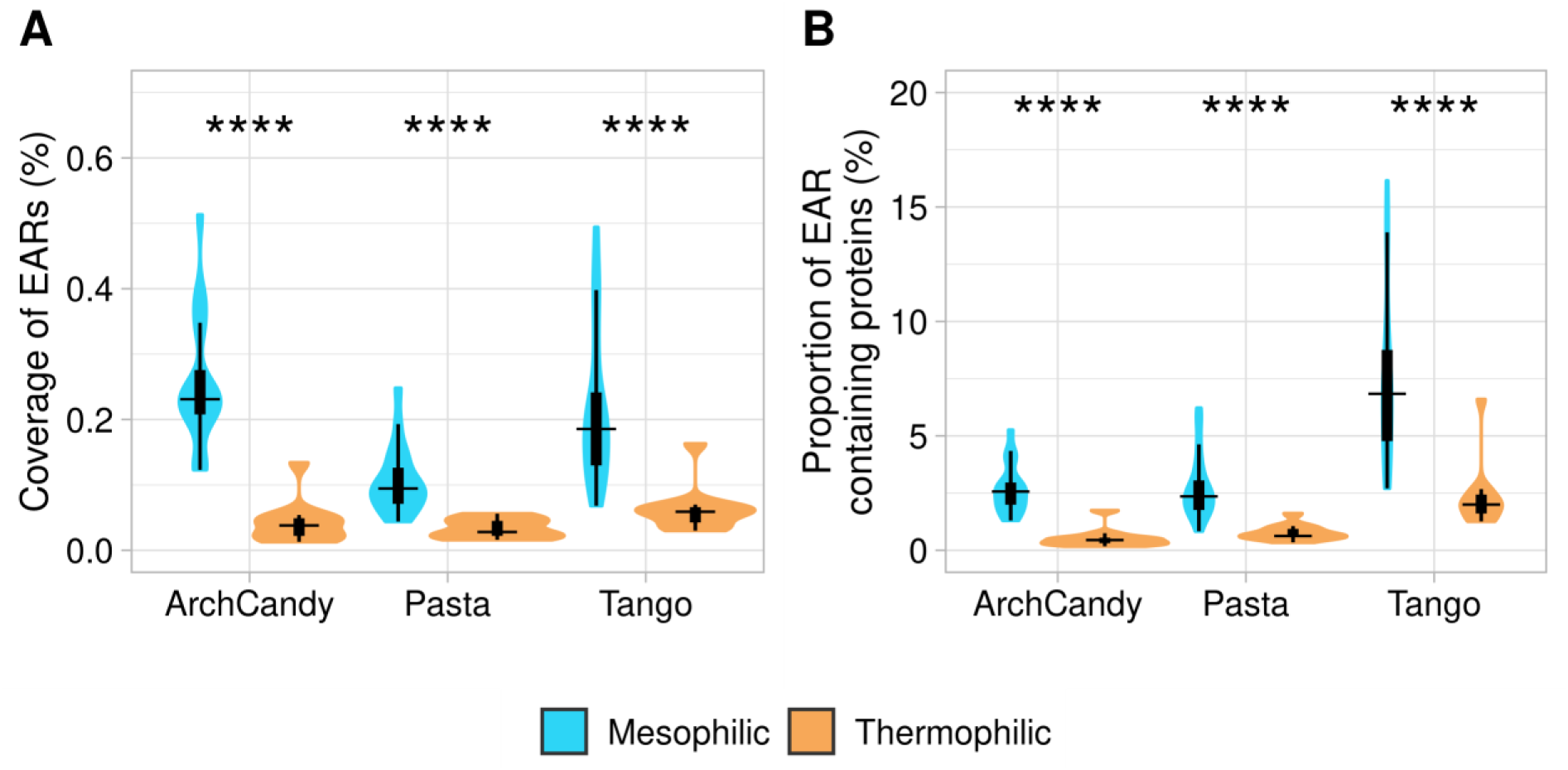
Coverage of EAR (A) and proportion of EAR-containing proteins (B) in mesophilic (blue) and thermophilic organisms (orange). The analyzed set of thermophilic organisms is: Chloroflexus aurantiacus, Thermodesulfovibrio yellowstonii, Dictyoglomus turgidum, Nanoarchaeum equitans, Sulfolobus solfataricus, Thermotoga maritima, Archaeoglobus fulgidus, Thermococcus kodakaraensis, Methanocaldococcus jannaschii, Candidatus korarchaeum, Aquifex aeolicus. For statistical analysis between mesophilic and thermophilic organisms we performed a t-test for predictors individually (ns: non-significant; *: p-value < 0.05; **: p-value < 0.01; ***: p-value < 0.001; ****: p-value < 0.0001).

### Occurrence of EARs in proteins depending on their length

In general, the longer the protein chain, the higher the probability for it to have both ARs and EARs. One would expect that if the ARs or EARs are uniformly distributed in protein sequences, their occurrence would correlate linearly with protein length. To see the tendency better, one can normalize the occurrence of ARs/EARs by dividing it by protein length. Previously, similar analyses have been done for the ARs using bacterial proteins (25) and the human proteome (27). Both studies showed that the aggregation potential of a protein normalized by its length goes down with the increase of protein size. To compare this conclusion with our results from the 76 selected proteomes, we analyzed the normalized proportion of AR-containing proteins and normalized AR-coverage depending on length (Figure 5 A, C). In agreement with the previous studies, we observed a decrease in the normalized proportion of AR-containing proteins and AR coverage with length. The steady decrease starts after 500 residues. The graph of AR coverage has a sharp peak at around 350-residue length. Clustering proteins by MMseqs2 (51) at 30 % of sequence identity, we found that this peak contains a significant excess of G protein-coupled receptors having high AR coverage, explaining this anomaly. The 200-500 residue region with the highest AR coverage and proportion coincides with the length ranges where proteins are predicted to be the most structured (Figure 5E) and in general, it negatively correlates with the IDR coverage by protein length. Thus, the AR proportion and coverage curves can be explained by the fact that structured regions have a higher probability of containing ARs, and proteins of less than 500 residues are mostly structured.

**Figure 5.**
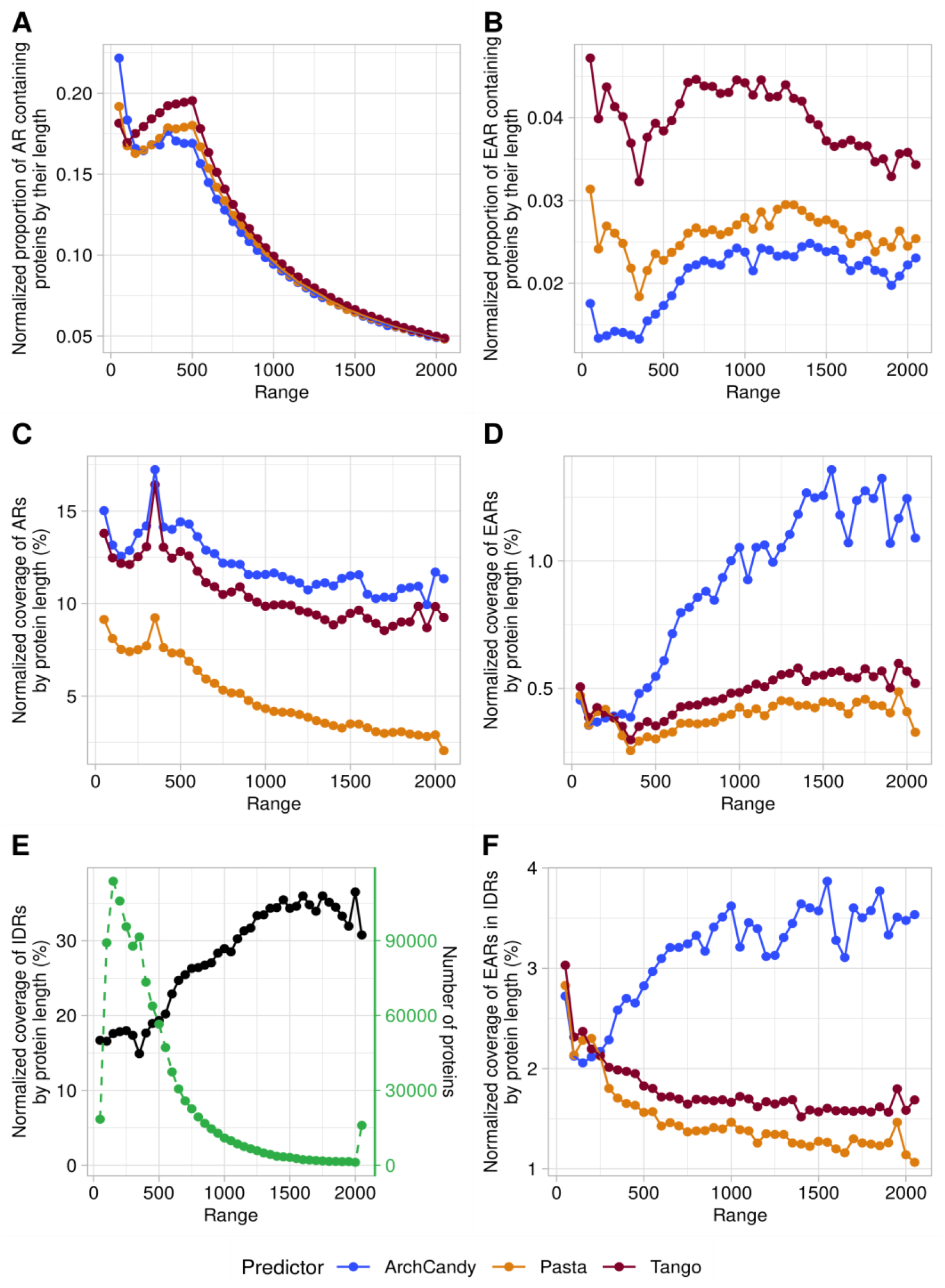
Plots of the proportion of AR (A) and EAR (B) containing proteins depending on the protein length. Plots of coverage of AR (C) and EAR (D) in proteins according to their length. Plots of coverage of IDR (E) and EAR in IDR (F). Proteins are grouped by subsets of 50 residues (e.g. 1-50, 51-100 etc). Proteins longer than 2000 were grouped into one subset. The predictors used have systematic biases at the terminal regions of proteins and this affects results on the short sequence lengths. To take this bias into account, we also run the predictors against a set of randomized sequences. This set contains proteins from our database with each sequence computationally shuffled, respecting the average amino acid composition of our database and having the same distribution of protein lengths. This allowed us to determine a correction coefficient which was used to adjust the values of EAR, AR and IDR.

The dependence of EARs on protein length demonstrates that it differs from ARs (Figure 5 B, D).

The predictors show a plateau with the lowest EAR-coverage for the shortest proteins (less than 350 residues), which steadily goes up for longer proteins. A similar trend is observed when we plot the dependence of the proportion of EAR-containing proteins by length.

The dependence of the coverage of IDRs against protein length (Figure 5E) is similar to the one of EARs, explaining the low aggregation potential of the short sequences by their tendency to be structured. Indeed, the region of 200-400 residues, which corresponds to the stable structural domains of proteins, has the lowest coverage of IDRs and EARs.

To see the tendency linked only to the characteristics inherent in the IDR sequences, we analyzed the dependence between the EAR coverage in IDRs and the length of proteins. The analysis shows that for TANGO and PASTA 2.0, shorter sequences have higher EAR coverage in IDRs. In contrast, ArchCandy 2.0 predicted an increase of EAR coverage in IDRs with protein length (Figure 5F). One explanation of this discrepancy between the predictors maybe the fact that ArchCandy predicts Asn/Gln-rich regions, which are frequently found in long proteins, as aggregation prone, while TANGO and PASTA do not.

Thus, we do not observe a decrease in aggregation potential with an increase of protein size when we consider EARs. The longer a protein chain, the higher its propensity to aggregate. Therefore, the question arises as to the mechanism preventing fibril formation of long proteins. One possible explanation can be that long proteins, having multiple IDRs, represent “steric brushes”, preventing their intermolecular interactions and aggregation due to the high entropic barrier (52).

### Occurrence of EAR-containing proteins in different cellular compartments

Proteins having different cellular localizations may differ in their aggregation potential. Therefore, we analyzed the occurrence of AR- and EAR-containing proteins in 4 major subcellular localizations: secreted proteins identified by SignalP (40), transmembrane proteins by using TMHMM (39), nuclear proteins with NLS (nuclear localization signals) found by SLiMs (41), and the remaining proteins that were considered mostly cytosolic (Figure 6). We observed similar levels of AR-containing proteins in all compartments except the transmembrane proteins, which have significantly higher levels (Supplementary Figure 2). The high level of AR-containing proteins among the transmembrane proteins was expected because their hydrophobic TM helices are detected as ARs by all predictors. The most striking observation was the high level of EAR-containing proteins in nucleus of eukaryotes, which is at least two times higher than in the other cellular localizations (Figure 6A). In line with this result, it has been shown previously that under stress conditions, proteins in the nucleus tend to form aggregates (53).

**Figure 6.**
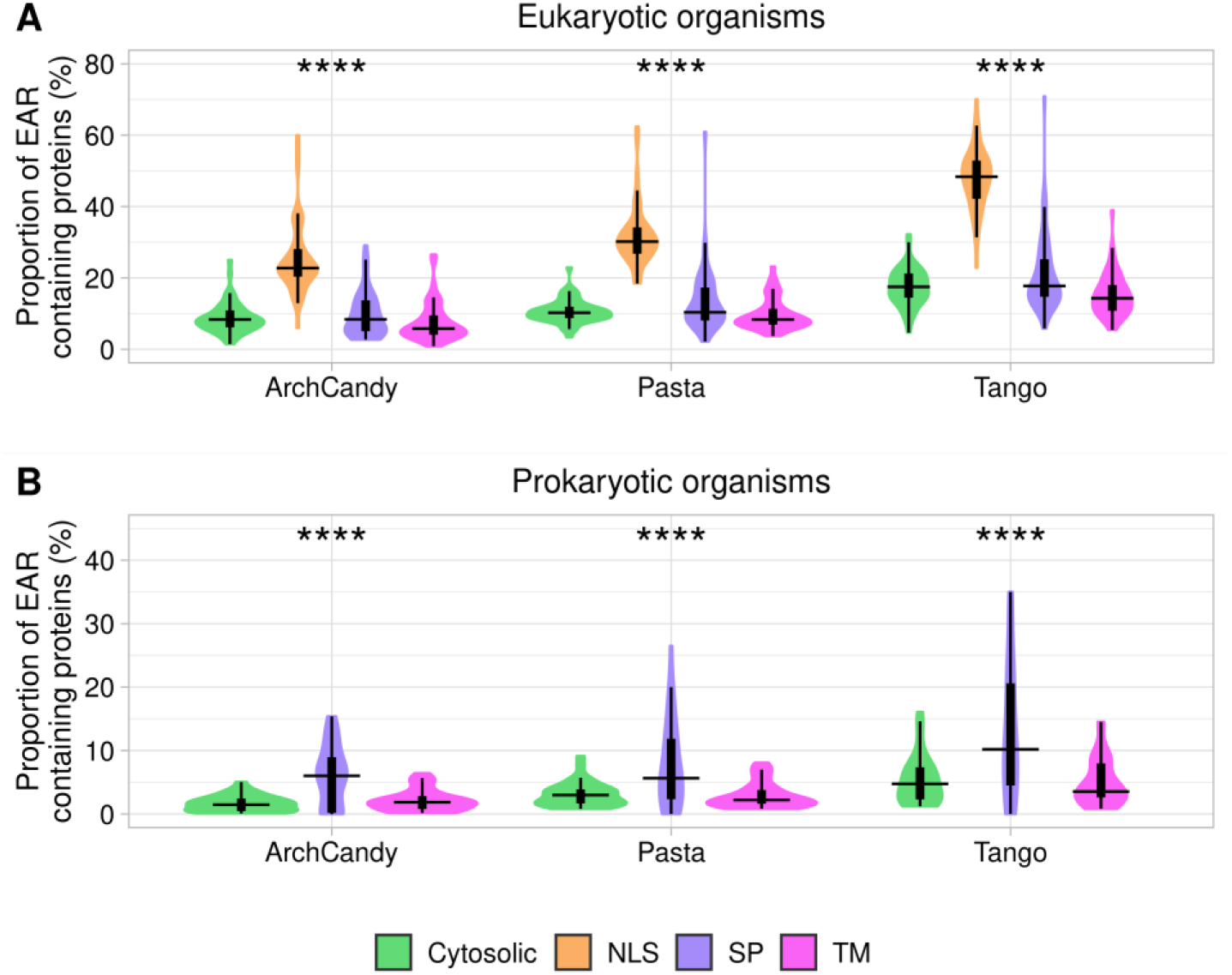
Plots of proportion of EAR-containing proteins according to the protein localization in eukaryotic (A) and prokaryotic (B) organisms. Proteins are splitted in four groups: cytosolic proteins, extracellular ones with signal peptides (SP), transmembrane proteins (TM) and nuclear proteins having nuclear localization signals (NLS). The proportions of the analyzed proteins are: in eukaryotes, 58.5% are cytosolic, 6.2% have SPs, 21.4% are transmembrane proteins and 13.9% have NLS. In prokaryotes, 71.9 % are cytosolic, 4.3% have SPs and 23.7% are transmembrane proteins. For statistical analysis between the different cell compartments we performed an anova test for the predictors individually (ns: nonsignificant; *: p-value < 0.05; **: p-value < 0.01; ***: p-value < 0.001; ****: p-value < 0.0001).

In prokaryotes, we observed more EAR-containing proteins among those involved in the secretory pathway in comparison to those present in the transmembrane and cytosol (Figure 6B). This tendency suggests that the secreted proteins being outside of the cell are under a reduced evolutionary pressure to avoid aggregation. Formation of aggregates out of the cell may be less deleterious for unicellular prokaryotic organisms in comparison with most of the eukaryotes, which can accumulate unwanted deposits within the extracellular space of their tissues. Moreover, it is known that many prokaryotes use secreted proteins to form functional amyloids (5).

### Relationship between cellular abundance of proteins and ARs/EARs frequencies

The amount of genome-wide data on gene expression has drastically increased in the past few years (54). The data comes from various technologies, organisms, and tissues (normal or disease related), making it difficult to compare them in a large-scale analysis. In this case, we find that the data from the Protein Abundance Database (PaxDb) (55) are the most suitable for our purposes. PaxDb represents protein abundance by “protein per million” (ppm) and by doing so, overcomes the problem of variability in cell size or dilutions in the samples used, making comparisons between them possible. The PaxDb has the protein expression level in different tissues and organs of organisms. Additionally, it provides the average abundance of a protein in the whole organism. We used this average abundance value to analyze the expression level of AR-/EAR-containing proteins, which are available both in PaxDb and in our dataset. Expression levels range from almost zero up to more than 100 000 ppm. The majority of proteins have values of less than 2 ppm. The number of proteins with the abundance more than 50 ppm drops significantly (Figure 7). Therefore, we grouped these proteins together in our analysis.

**Figure 7.**
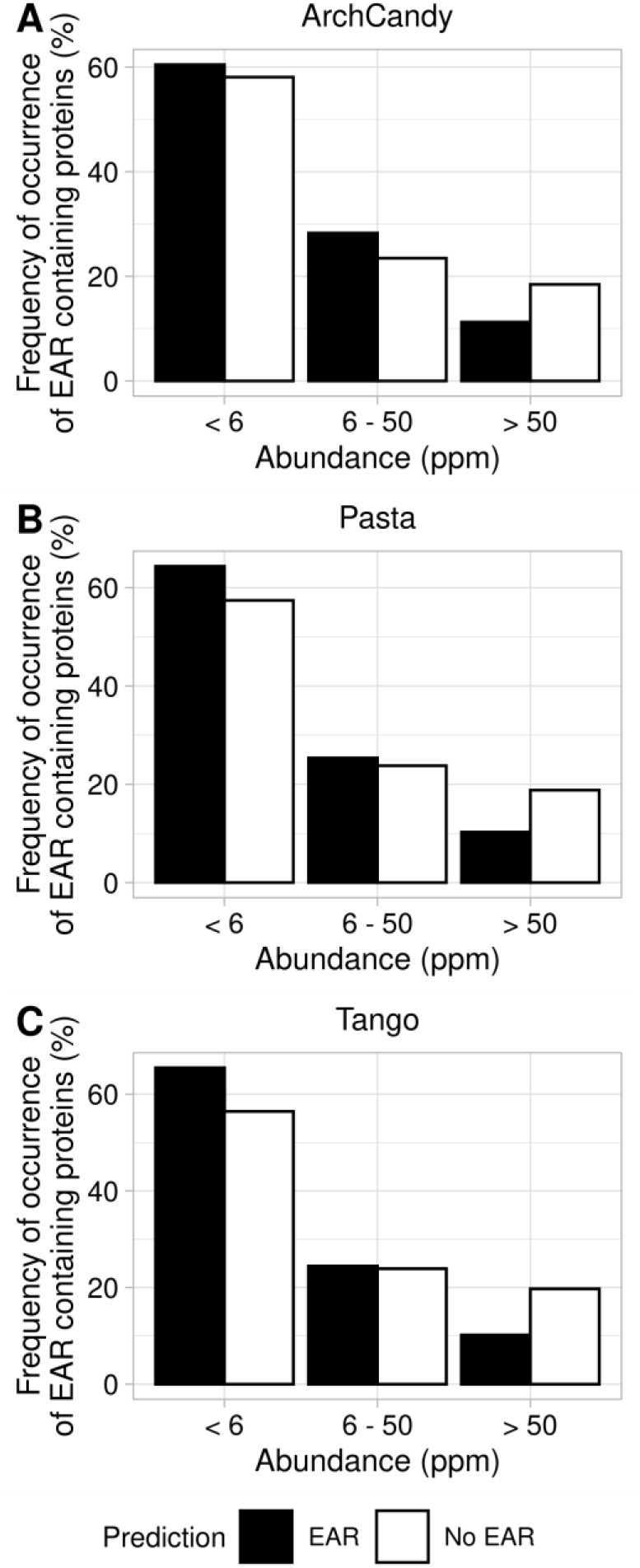
Frequency of occurrence of EAR -containing proteins predicted by ArchCandy 2.0 (A), Pasta 2.0 (B) and TANGO (C). Proteins are grouped based on their abundance in three groups: less than 5 ppm, 5-50 ppm, and more than 50 ppm.

The analysis revealed that the frequency of occurrence of EAR-containing proteins decreases with the ppm growth and is becoming lower than non-EAR-containing proteins. From the observed dependence of the difference between EAR- and non-EAR-containing proteins, we can conclude that highly expressed proteins are less prone to aggregate, with this finding being consistent in the three predictors used. We observed a similar tendency with the frequency of occurrence of AR-containing proteins depending on the abundance (see Supplementary Figure 3). It suggests that highly expressed proteins are under a greater selective pressure to avoid aggregation. This conclusion is in agreement with a previous study of human proteins also suggesting that aggregation-prone proteins and gene level expression are inversely correlated (56).

### EAR levels in essential proteins

As demonstrated previously, essential genes are subject to a greater selection pressure than non-essential genes (55, 57). It has also been shown that essential proteins are less prone to aggregation (20, 23). In order to find essential proteins in our database, we used the DEG database of essential proteins (58) and run BLAST program with E-value < 0,001 (59). By this approach we identified 705692 essential proteins (~62,6 %) in our database. Analysis of these proteins by the three predictors showed a lower EAR coverage, and in a lesser extent, proportion of essential and non-essential EAR-containing proteins in eukaryotes (Figure 8 A, C). Our results are in agreement with previous conclusions that essential proteins have a lower aggregation score than non-essential proteins (23). In prokaryotes, we observe the opposite tendency (Figure 8B, D). Previous analyses of ARs (not EARs) made on a smaller scale in bacteria (25) have shown that essential proteins have less ARs. Our results of the AR analysis in prokaryotes (see Supplementary Figure 4) are in agreement with this conclusion.

**Figure 8.**
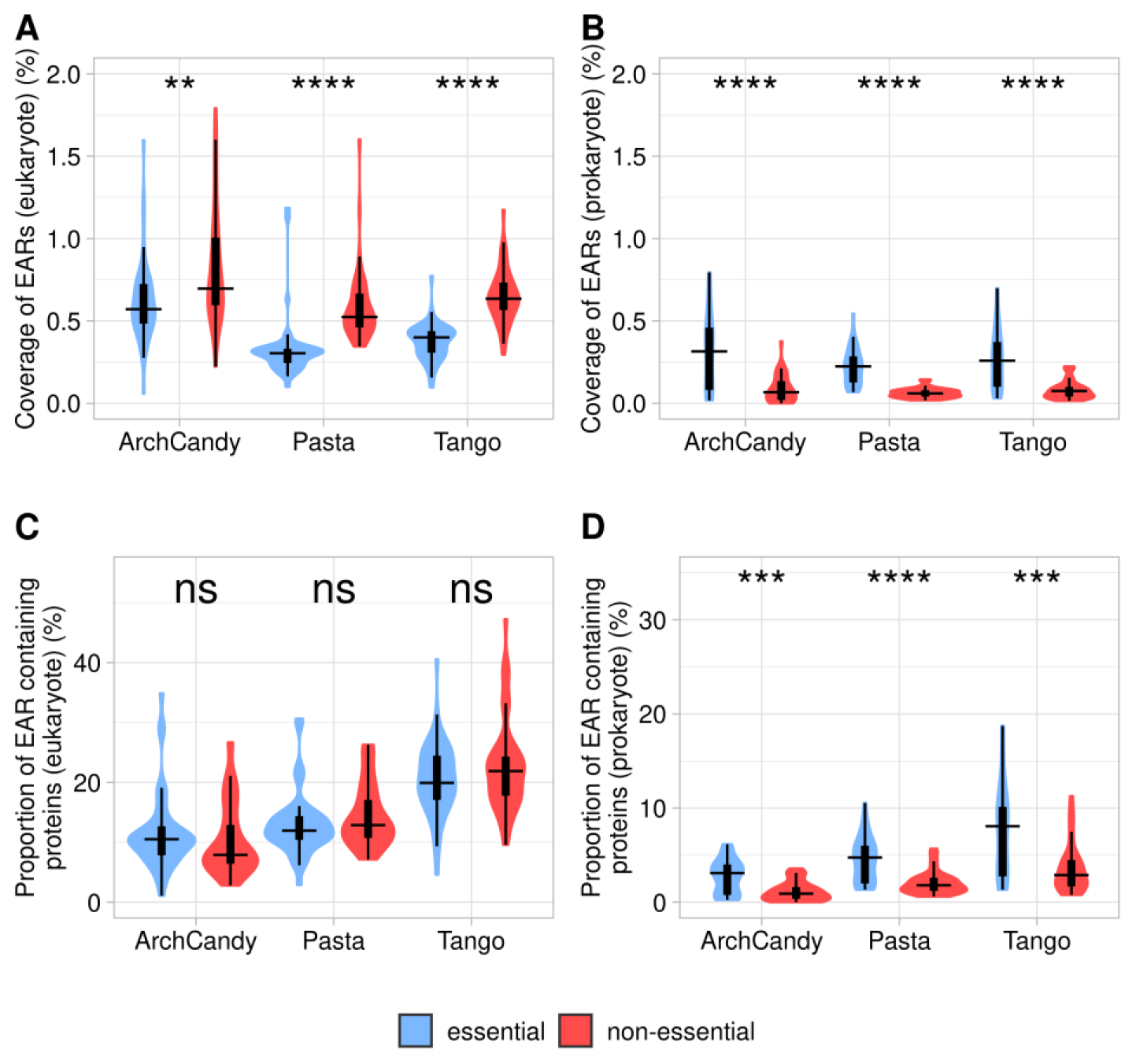
Coverage of EARs in essential and non-essential proteins in eukaryote (A) and prokaryote (B) organisms. Proportion of EAR-containing proteins known as essential or non-essential in eukaryote (C) and prokaryote (D) organisms. For statistical analysis between essential and non-essential proteins we performed a t-test for amyloidogenic predictors individually (ns: non-significant; *: p-value < 0.05; **: p-value < 0.01; ***: p-value < 0.001; ****: p-value < 0.0001).

### Short Linear Motifs (SLiMs) in EARs

A significant portion of protein interactions are mediated by short linear motifs (SLiMs) preferentially found in IDRs (41). As EARs are also located within the IDRs, it was interesting to analyze the co-occurrence of SLiMs and EARs in proteins. Although both prokaryotes and eukaryotes have functional SLiMs, the eukaryotic linear motifs are more common, as well as better classified and documented. Most of the eukaryotic SLiMs can be found in the ELM resource (41) alongside with their descriptions, experimental evidence from the literature, and Regular Expressions (RegEx) of the recurrent patterns. Therefore, we focused our analysis on the SLiMs from eukaryotes. For this purpose, we applied the RegEx from the ELM database (41) to the IDRs and EARs determined by our pipeline (33). The SLiMs are subdivided into 6 major classes: (LIG) ligand binding motifs and (DOC) docking sites both involved in protein-protein interactions of the functional complexes, (MOD) modification sites covering several post-translational modifications of proteins (e.g. phosphorylation, palmitoylation, glucosylation), (DEG) sites of proteins that are important in regulation of protein degradation rates, (TRG) targeting sites responsible for protein sorting in cellular compartments and (CLV) specific cleavage sites.

The results of all three aggregation predictors showed that EAR-containing proteins are enriched in SLiMs in comparison to IDR-containing proteins without EARs (Figure 9). By using the exact Fisher test, we were able to select SLiMs, which are significantly enriched in EAR-containing proteins compared to IDR-containing proteins alone. Interestingly, 20 of the 25 degradation motifs (proteasome pathway) from DEG class occur more frequently in EAR-containing proteins than in non-EAR-containing proteins (Figure 9). 17 of the 22 TRGs are also more frequently present in EAR-containing proteins than in IDR-containing proteins. Among them, 3 SLiMs were found to be Endosome-Lysosome-Basolateral sorting signals. These results suggest that EAR-containing proteins may be more susceptible to being degraded by proteasome and lysosome pathways compared to just IDR-containing proteins. This might be a strategy used by organisms to prevent protein aggregation by increasing the degradation of potential aggregation-prone proteins. Cleavage sites (CLV) are less prevalent in EARs, which may prevent the release of smaller amyloidogenic peptides such as the well-known Aβ-peptide (60).

**Figure 9.**
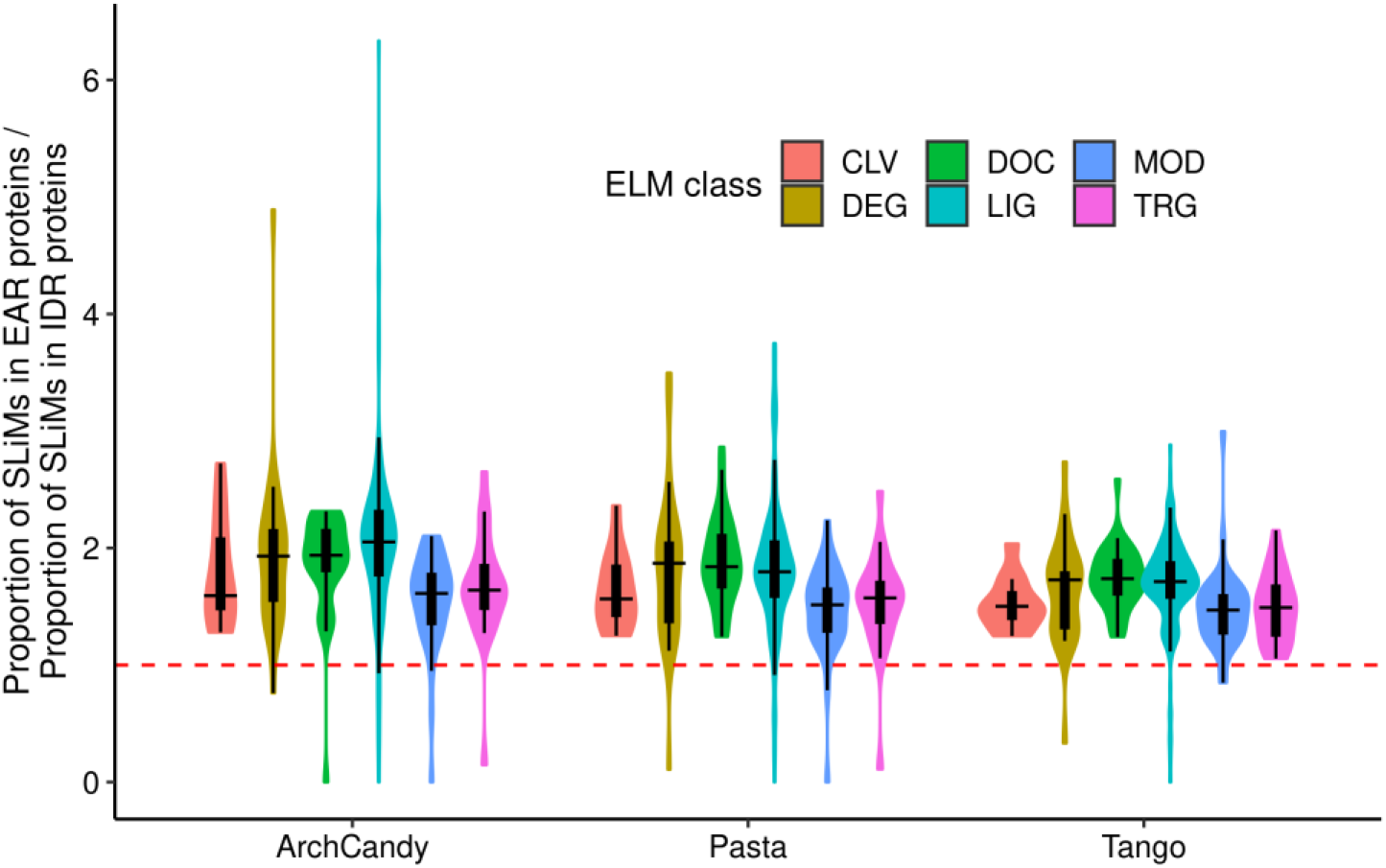
Ratio of proportions of SLiMs in EAR-containing proteins and in IDR-containing proteins without EAR, predicted by three predictors. Each dot represents a given SLiMs grouped in 6 classes denoted by different colors. The majority of the SLiMs have their ratios greater than 1.0 (red dotted line), meaning that they are enriched in EAR-containing proteins.

### Functional domains enriched in EAR-containing proteins

With a method similar to the SliMs enrichment, we tried to identify functional Pfam domains enriched in EAR-containing proteins. Of the 15116 known Pfam domains, only 1410 are significantly more prevalent in EAR-containing proteins predicted by ArchCandy 2.0 (exact Fisher test, p-value < 0.001). 484 of them belong to 154 clans according to the classification of Pfam. The functional domains and clans that came on top are: nucleoporin FG repeat region (CL0647), RNA recognition motif domains (CL0221) and zinc-finger domains (CL0511, CL0390) (see Supplementary data 1). We searched the experimental evidence of aggregation by these domains in the literature and found that the nucleoporin proteins are known to form amyloids (61). EAR-containing proteins predicted by both ArchCandy 2.0 and TANGO are positively enriched in nucleoporin FG repeat region (CL0647). From the known functional amyloids described in the literature, we also found back RIPK1 and RIPK3 (8, 9) and PMEL17 (62), which were conserved in 6 distinct proteins from mammalians with the prediction of ArchCandy 2.0 but not Pasta 2.0 or TANGO.

Previous studies of Pfam domains and gene ontology (GO) term enrichment in amyloidogenic proteins (24, 28) pointed out the over-representation of membrane transport activity, pH and ion regulation and even cytoskeleton organization. However, they considered ARs not EARs. Therefore, we did not find most of the afore mentioned functions in our analysis.

### Conservation of EARs sequences

Another approach to find new functional amyloids is to search for EARs that are conserved among different species. For this purpose, we reduced EARs predicted by either ArchCandy 2.0, TANGO or Pasta 2.0 with CD-HIT (63) at 70 % sequence identity and 90 % of coverage, to obtain a non-redundant set of the EARs (Table 1). Then, we ran BLAST (59) for each EAR sequence against all proteins from our redundant database to select only conserved EAR sequences (Figure 10). This gave us for each EAR a Multiple Sequence Alignment (MSA) of similar sequences found in other proteins. Some sequences of the MSA were EARs and the others were not according to the predictors. We selected the MSAs with EARs in more than five other proteins and further reduced the MSA number by merging those that shared at least 80 % of the same conserved EAR. This clustering results found 2218, 869 and 178 of the most conserved EAR sequences for ArchCandy 2.0, Pasta 2.0 and TANGO, respectively (Table 1). We observed that only a small number of EAR sequences are conserved out of more than one million proteins. Among them we found already known functional amyloids, such as RIPK3 and RIPK1 (8) and PMEL17 (62). This suggests that the list of conserved EARs found by this protocol (Supplementary data 2) can be used for detection and experimental tests of new functional amyloids.

**Table 1.** Number of EARs at each step of the protocol.

| Predictor | Number of non-redundant EARs | EARs found one time in MSA | EARs found 2 to 5 times in MSA | EARs found more than 5 times in MSA | Number of clusters with the most conserved EARs |
| --- | --- | --- | --- | --- | --- |
| ArchCandy 2.0 | 93229 | 72153 (77.4%) | 16124 (17.3%) | 4952 (5.3%) | 2218 |
| Pasta 2.0 | 42997 | 35412 (82.4%) | 5683 (13.2%) | 1902 (4.4%) | 869 |
| TANGO | 13816 | 12342 (89.3%) | 1219 (8.8%) | 255 (1.8%) | 178 |

**Figure 10.**
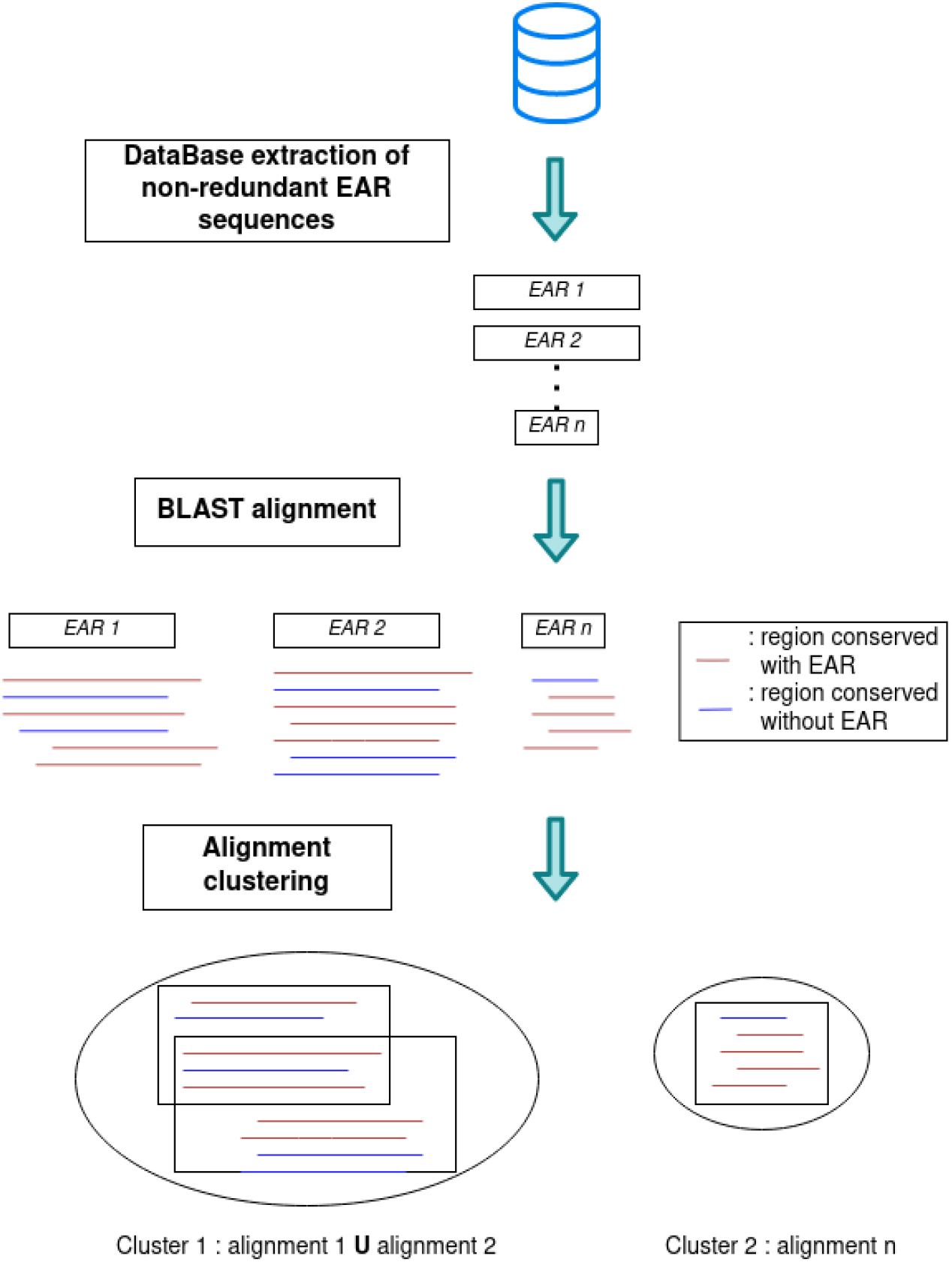
Protocol for the evaluation of EAR sequence conservation.

## DISCUSSION

The recent progress with computational approaches predicting aggregation (10, 12, 13, 15, 16, 18, 31, 33), and an increasing number of whole-proteome sequencing data, opened an avenue for the comprehensive census of aggregation-prone regions in proteins. In this work, we performed the detailed analysis of 76 full reference proteomes from the UniProt databank. As a result, a number of interesting correlations, confirmed by all the predictors used in this work (ArchCandy 2.0, Pasta 2.0 and TANGO), were discovered. First, we detected a significantly lower percentage of EAR-containing proteins (about 10%) in comparison with a high percentage of AR-containing proteins in proteomes (about 80%). The number of EARs correlates better with a small number of the known proteins forming aggregates *in vivo*, and, therefore, EARs can be suggested as a more precise measure of the aggregation potential of proteins. We showed that there are more ARs in prokaryotes than in eukaryotes and that this tendency is inverted for EARs. Second, we found that the thermophilic prokaryotes have significantly less EARs and ARs in comparison to mesophilic prokaryotes. The correlation may reflect an evolutionary pressure on the thermophilic proteins, because the amyloid formation rate constant increases with temperature (50).

Additionally, in agreement with previous studies, we observed small decrease in the normalized AR coverage with protein length. However, we do not observe a decrease in the aggregation potential of sequences with an increase of protein size when we consider EARs. In our opinion, the mechanism of prevention of aggregation of long proteins has an entropic basis, where the other parts of the chain generate repulsive forces for intermolecular interactions similar to molecular brushes.

It worth mentioning that our analysis did not confirm previously published conclusions that the average aggregation propensity of a proteome correlates inversely with the complexity and longevity of the studied organisms (29).

It was also shown that proteins having different cellular localizations differ in the aggregation potential. For example, the level of EAR-containing proteins in nuclear proteins of eukaryotes is about two times higher than in the other cellular localizations. In prokaryotes, we observed more EAR-containing proteins among those involved in the secretory pathway in comparison to the transmembrane and cytosolic proteins. This tendency suggests that the secreted proteins being outside of the cell are under a reduced evolutionary pressure to avoid aggregation. Remarkably, a great majority of EAR-containing proteins are enriched in SLiMs in comparison to IDR-containing proteins without EARs. We also noticed that highly expressed proteins are less prone to aggregate suggesting that highly expressed proteins are under a greater negative selective pressure in order to avoid the aggregation. Finally, we revealed a greater level of aggregation predicted in non-essential proteins compared to essential proteins.

Thus, we performed the census of the aggregation-prone regions in proteomes. A number of new relationships found in this work led us to a better understanding of the functional and evolutionary relations of protein aggregation in organisms from the three kingdoms of life: eukaryote, bacteria and archaea. Beyond this, our study opens up new opportunities for a number of experimental tests.

## Supporting information

List of species

## ACKNOWLEDGMENT

The authors thank Priya Amin for her assistance with the English.

## FUNDING

This work was supported by REFRACT project with Latin America in RISE program (2018-2023) H2020-MSCA-RISE-2018 to A.V.K.; Azerbaijan National Academy of Sciences and The Ministry of Science and Education of Azerbaijan to Z.O.; the Ministère de l’Education Nationale de la Recherche et de Technologie (MENRT) to E.V. and F.R.; by a CNRS PhD fellowship to T.F.

## Author Contributions

Conceptualization, A.V.K., T.F. and E.V.; methodology, A.V.K., T.F. and E.V.; software, T.F. and E.V.; data curation, T.F., F.R. and Z.O.; writing—original draft preparation, T.F. and A.V.K.; writing—review and editing, A.V.K., T.F., Z.O, F.R, E.V.; supervision, A.V.K.

## CONFLICT OF INTEREST

Authors declare no conflict of interest.

## SUPPLEMENTARY DATA

**Supplementary Figure 1.**
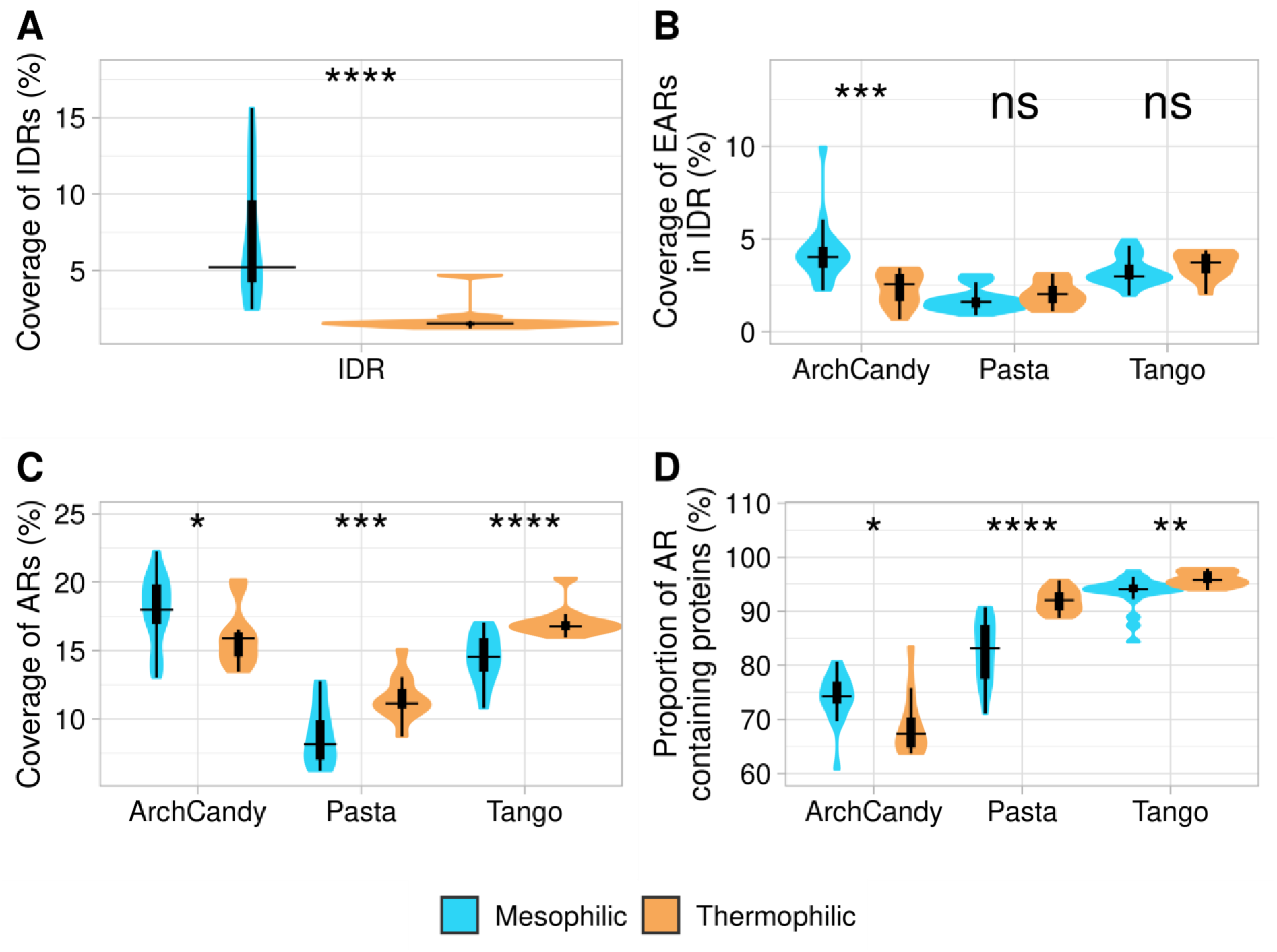
Coverage of IDR (A), EAR in IDR (B), AR (C) and proportion of AR containing proteins (D) in mesophilic (blue) and thermophilic organisms (red). The analyzed set of thermophilic organisms is: Chloroflexus aurantiacus, Thermodesulfovibrio yellowstonii, Dictyoglomus turgidum, Nanoarchaeum equitans, Sulfolobus solfataricus, Thermotoga maritima, Archaeoglobus fulgidus, Thermococcus kodakaraensis, Methanocaldococcus jannaschii, Candidatus korarchaeum, Aquifex aeolicus.

**Supplementary Figure 2.**
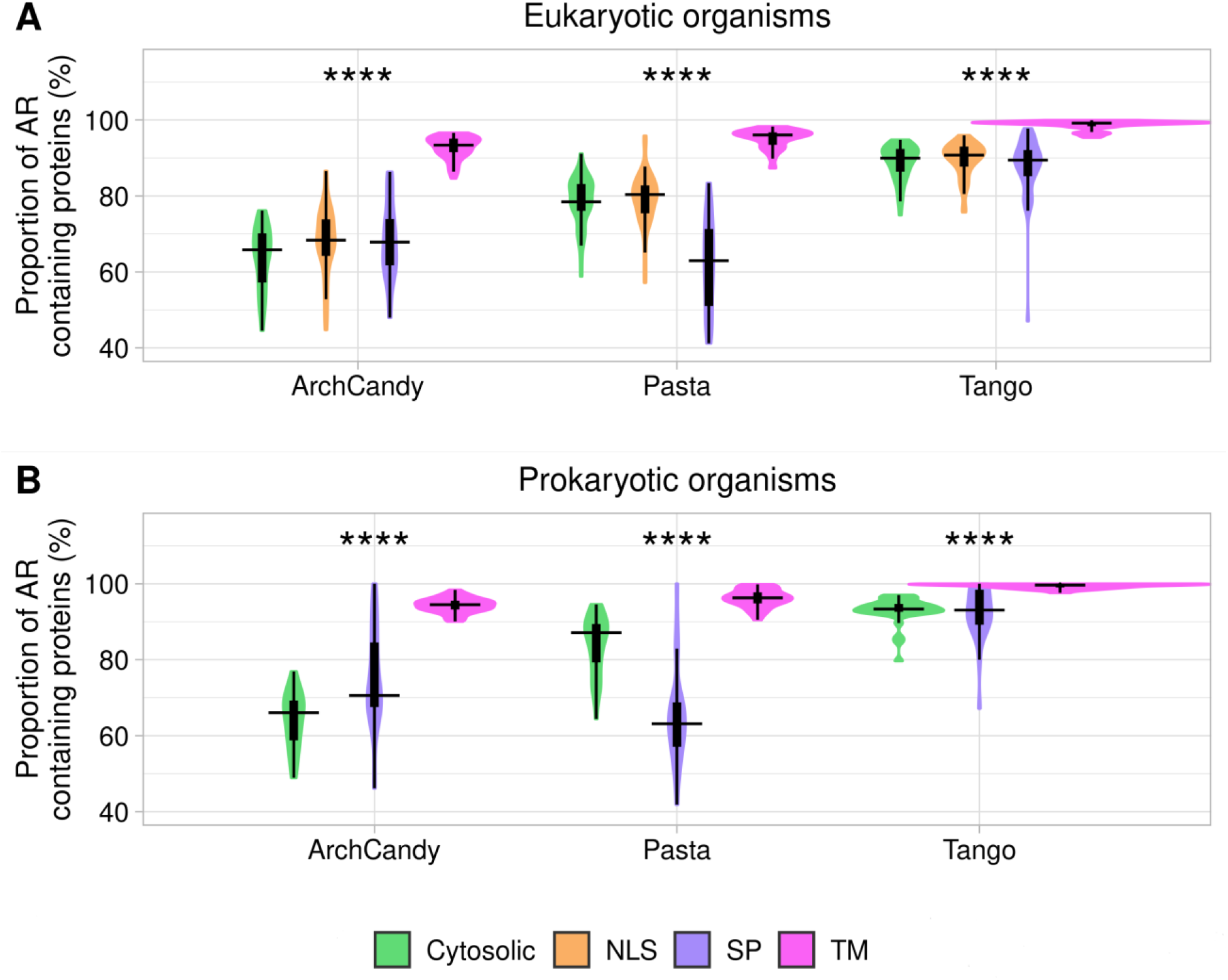
Plots of proportion of AR containing proteins according to the protein localization in eukaryotic (A) and prokaryotic (B) organisms. Proteins are splitted in four groups: cytosolic proteins, extracellular ones with signal peptides (SP), transmembrane proteins (TM) and nuclear proteins having nuclear localization signals (NLS). The proportions of the analyzed proteins are: in eukaryotes, 58.5% are cytosolic, 6.2% have SPs, 21.4% are transmembrane proteins and 13.9% have NLS. In prokaryotes, 71.9 % are cytosolic, 4.3% have SPs and 23.7% are transmembrane proteins

**Supplementary Figure 3.**
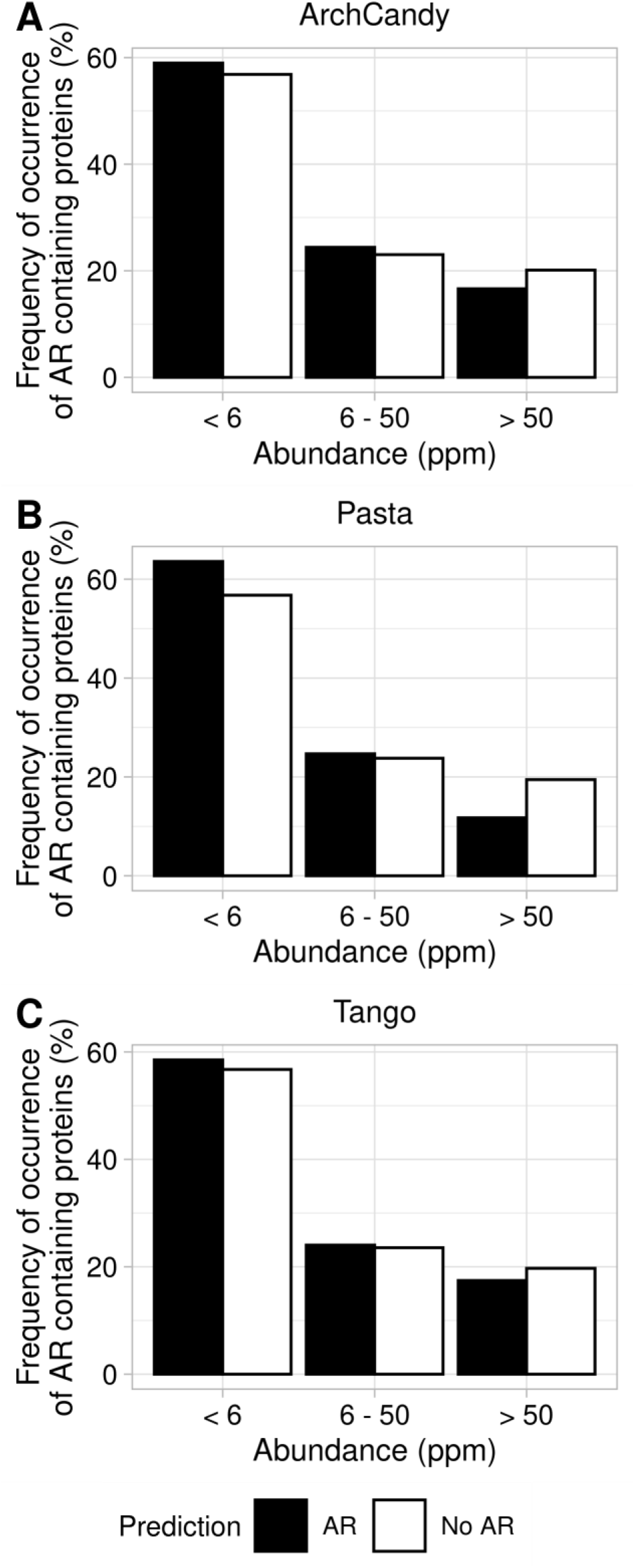
Frequency of occurrence of AR containing proteins predicted by ArchCandy 2.0 (A), Pasta 2.0 (B) and TANGO (C). Proteins are grouped based on their abundance in ranges of 5 ppm, proteins with 50 ppm or more are grouped in one range.

**Supplementary Figure 4.**
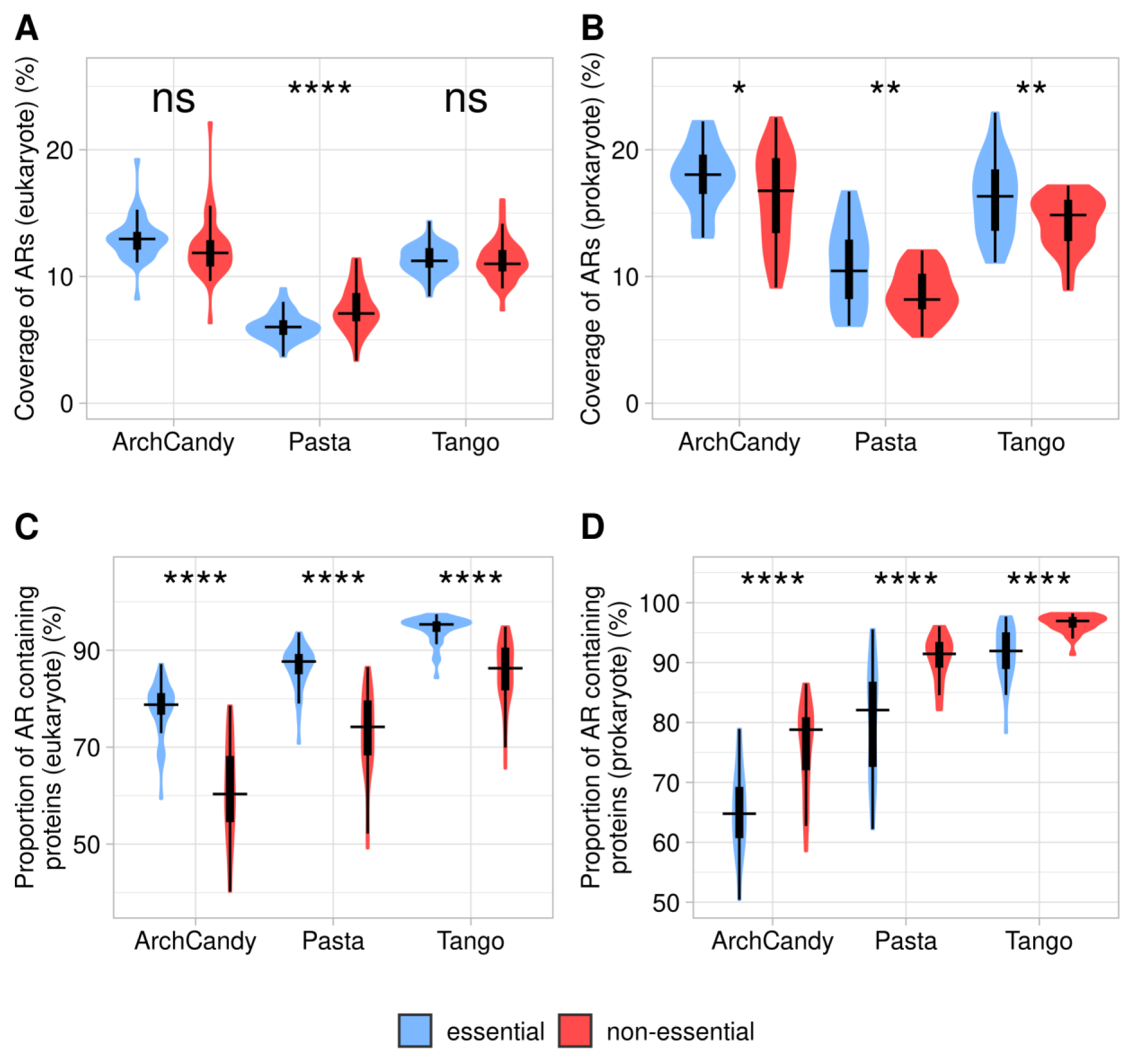
Coverage of AR in essential and non-essential proteins in eukaryote (A) and prokaryote (B) organisms. Proportion of AR containing proteins known as essential or non-essential in eukaryote (C) and prokaryote (D) organisms.

## Notes

### Competing Interest Statement

The authors have declared no competing interest.

